# Sensory Neural Noise as a Limiting Factor in Visual Working Memory Precision in Neurotypicals and Individuals with Schizophrenia

**DOI:** 10.1101/2025.10.20.683426

**Authors:** Woon Ju Park, Megan Ichinose, Gengshi Hu, Geoffrey F. Woodman, Duje Tadin, Sohee Park

## Abstract

The neural mechanisms that determine the capacity limits of working memory (WM) are not well understood, with traditional views identifying prefrontal circuitry as the main source of WM limits. Sensory noise, however, remains an underexplored explanation for WM limits despite its unavoidable influence on brain systems that encode and relay sensory information. Here, we show that individual differences in internal sensory noise during visual processing predict visual WM capacity. We further demonstrate that the well-documented WM deficits observed in schizophrenia can be associated with atypically high levels of visual sensory noise, offering a new explanatory framework for these deficits. By experimentally manipulating sensory noise via transcranial direct current stimulation applied to the visual cortex, we also show that changes in sensory noise led to corresponding changes in visual WM precision in individuals with schizophrenia. Finally, a computational model demonstrates that the same sensory noise measured during visual perception can limit WM precision during WM maintenance. These findings show that sensory noise may explain a significant amount of variance in WM function in both neurotypical adults and individuals with schizophrenia.

## INTRODUCTION

Working memory (WM) is central to flexible and adaptive human cognition, allowing intelligent guidance of behavior ^1^. The efficacy of WM is determined by its capacity, with capacity limitations leading to impaired information processing ^2–4^. WM deficits are present in a wide range of neuropsychiatric conditions, including schizophrenia, anxiety disorders, and dementias ^5–9^, and have a substantial impact on real-world functioning ^10^. Thus, elucidating mechanisms that determine the capacity of WM is of broad scientific and clinical importance. WM storage involves coordinating the functioning of multiple sensory, attentional, and cognitive control mechanisms ^11–13^. Traditional theoretical approaches have focused on prefrontal access and control as the primary constraint on WM effectiveness ^14–16^, and it remains controversial whether sensory processing plays a key role in WM capacity limitations ^17–19^.

Limitations in sensory processing have been overlooked as a critical determinant of WM capacity largely because the fidelity of sensory processing far exceeds the WM capacity for sensory information ^20^. At first glance, this makes sensory processing an unlikely culprit for WM limitations. The argument, however, fails to consider how variations in noise levels within sensory processing could cascade and affect brain functions that rely on sensory processing ^21^. Noise is ubiquitous in neural processing ^22^: It is present in sensory transduction, cellular processes of neurons, neural responses, and motor outputs ^23^, limiting the efficiency of the brain’s communication channels ^24^. Noise present at earlier stages of sensory processing is also subject to amplification during typical gain processes ^21^. Thus, atypical increases in sensory noise may have unavoidable cascading effects on a wide range of brain functions that rely on sensory inputs, including WM.

Electrophysiological, neuroimaging, and brain stimulation studies demonstrate that sensory areas remain engaged during the maintenance of visual WM representations ^25–27^, forming the basis of the *sensory-recruitment hypothesis* ^17,21^. Building on this view, we test the *sensory-noise hypothesis*, focusing on how sensory noise may limit WM capacity. We reasoned that visual WM may be particularly and uniquely vulnerable to sensory noise because it depends on the fidelity of sensory signals both for its initial encoding and for the maintenance of stored sensory information. In this framework, visual WM processing may be affected by sensory noise both in the feed-forward processing of visual information to be stored in WM and in the active maintenance of that information.

This framework makes several important predictions. First, it predicts that WM capacity limits will be systematically related to sensory noise levels in the general population. Second, it suggests that atypically high sensory noise could contribute to WM deficits observed in neuropsychiatric populations, such as schizophrenia, where limitations are often attributed to higher-level cognitive dysfunctions ^28^. Third, the framework also predicts that these limits should be changeable if we could manipulate noise levels in the visual cortex itself. Finally, it predicts that sensory noise estimated during perception should propagate and be amplified during the maintenance stage, thereby limiting WM precision.

We tested these predictions using converging methods of psychophysics and computational modeling, brain stimulation, and clinical science. Psychophysics quantified sensory noise during visual perception, computational modeling formalized how this noise limits WM precision, and brain stimulation provided a causal means to manipulate sensory noise in visual cortex. In our clinical work, we focused on schizophrenia where WM deficits have been proposed as an endophenotypic marker and a core feature of this condition ^8,9^. WM deficits in schizophrenia have been characterized as a failure of high-level executive control ^29^. However, schizophrenia is also associated with broadly atypical sensory processing ^30–33^, with implications for increased neural noise ^34–36^. Together with our sensory noise hypothesis of WM, these suggest a yet untested possibility that WM deficits in schizophrenia may derive from impaired sensory mechanisms.

## RESULTS

To test whether sensory noise limits visual WM, we asked both neurotypical adults (NT) and individuals with schizophrenia or schizoaffective disorder (SZ) to complete visual WM and orientation discrimination tasks (Study 1; 57 participants, 30 NT, 27 SZ, Table 1). The visual WM task required participants to remember locations and orientations of one, two, or four gratings and reproduce the orientation of a probed item after a 1 second delay with an input dial (Figure 1A; Methods). WM recall variance was the primary performance metric of interest. Less precise WM performance is reflected by higher recall variance, as demonstrated by the characteristic increase in recall variance (i.e., decline in precision of the reported orientation) as the number of items remembered increases ^37,38^. Given that internal noise is commonly assumed to add variability to the internal representation ^21,39^, we hypothesized that elevated sensory noise would lead to higher WM recall variance.

**Table 1.**
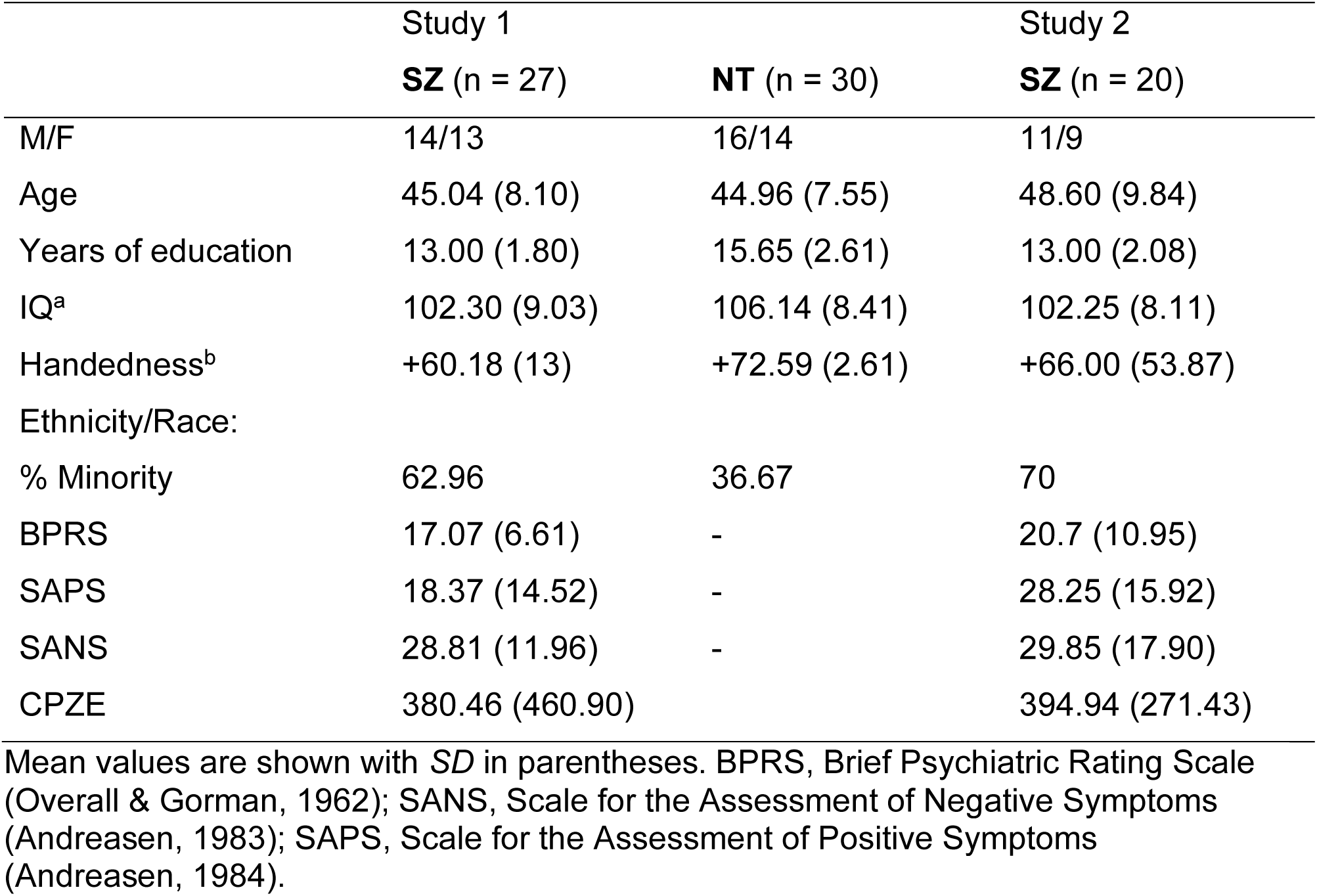
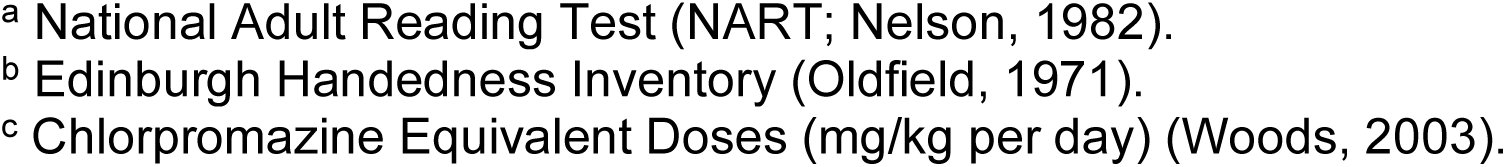
Demographic and Clinical Information for Studies 1 and 2.

|  | Study 1 |  | Study 2 |
| --- | --- | --- | --- |
|  | **SZ** (n = 27) | **NT** (n = 30) | **SZ** (n = 20) |
| M/F | 14/13 | 16/14 | 11/9 |
| Age | 45.04 (8.10) | 44.96 (7.55) | 48.60 (9.84) |
| Years of education | 13.00 (1.80) | 15.65 (2.61) | 13.00 (2.08) |
| IQ<sup>a</sup> | 102.30 (9.03) | 106.14 (8.41) | 102.25 (8.11) |
| Handedness<sup>b</sup> | +60.18 (13) | +72.59 (2.61) | +66.00 (53.87) |
| Ethnicity/Race: |  |  |  |
| % Minority | 62.96 | 36.67 | 70 |
| BPRS | 17.07 (6.61) | - | 20.7 (10.95) |
| SAPS | 18.37 (14.52) | - | 28.25 (15.92) |
| SANS | 28.81 (11.96) | - | 29.85 (17.90) |
| CPZE | 380.46 (460.90) |  | 394.94 (271.43) |

Mean values are shown with *SD* in parentheses. BPRS, Brief Psychiatric Rating Scale (Overall & Gorman, 1962); SANS, Scale for the Assessment of Negative Symptoms (Andreasen, 1983); SAPS, Scale for the Assessment of Positive Symptoms (Andreasen, 1984).
<sup>a</sup> National Adult Reading Test (NART; Nelson, 1982). <sup>b</sup> Edinburgh Handedness Inventory (Oldfield, 1971). <sup>c</sup> Chlorpromazine Equivalent Doses (mg/kg per day) (Woods, 2003).

**Figure 1.**
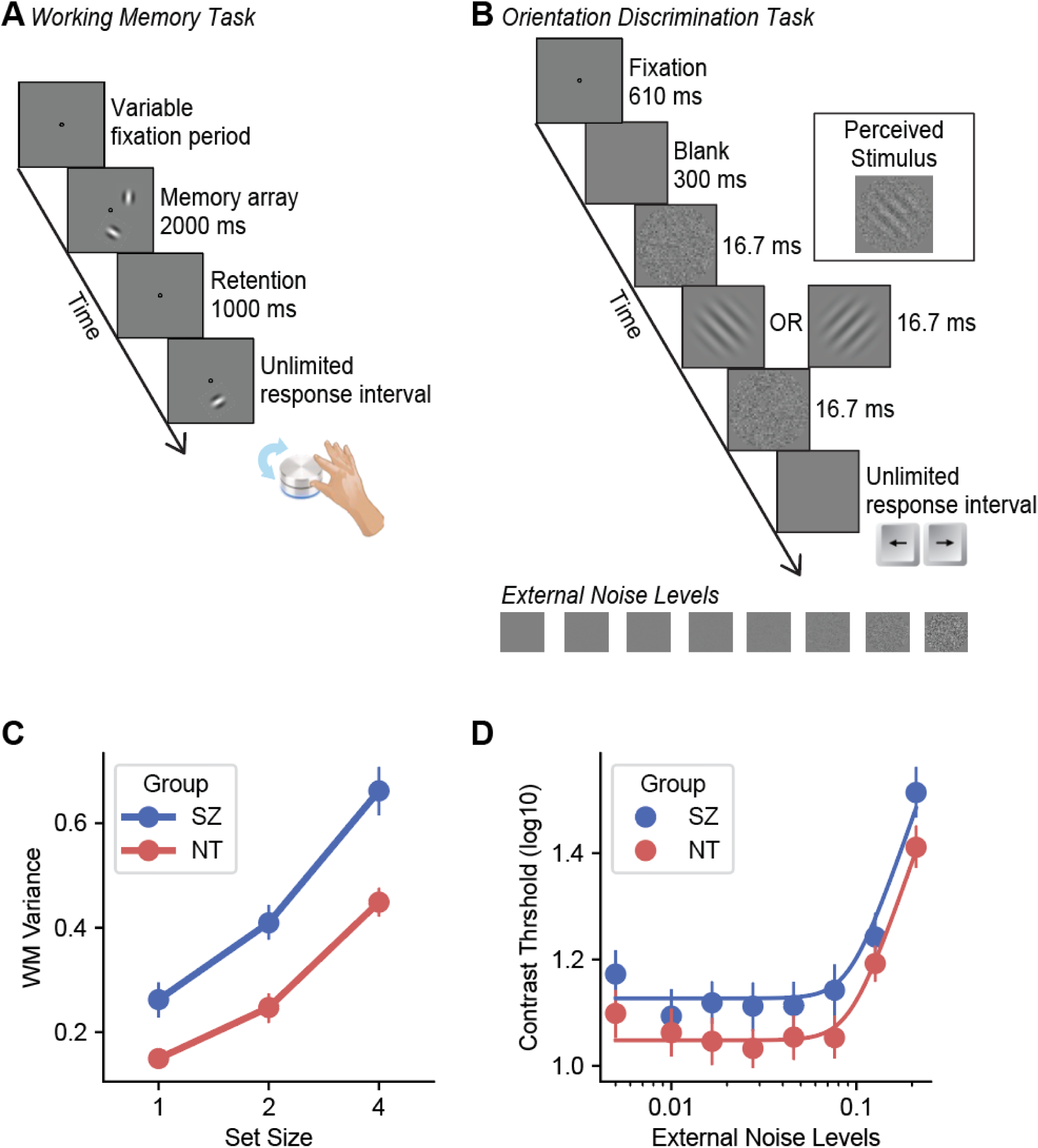
**A. & B.** Procedures for the visual WM task and the orientation discrimination task. See main text and Methods for details. **C.** Results for the visual WM task. Both NT (red) and SZ (blue) showed worse WM performance with increasing set size, with SZ performing significantly worse than NT. **D**. Results for the orientation discrimination task. Both NT (red) and SZ (blue) showed the characteristic non-linear increase in contrast thresholds with increasing external noise. SZ performed overall worse than NT, best explained by higher baseline internal noise and unfiltered external noise in SZ (relevant statistics are in the main text).

To quantitatively estimate sensory noise in visual processing, we used the equivalent noise paradigm—a well-established psychophysical approach that has been extensively validated in both basic vision science ^22,40,41^ and clinical populations ^42,43^. In this paradigm, participants discriminated coarse orientation differences (±45° tilt from vertical) of centrally presented gratings (1 cycle/°) embedded in varying levels of external noise (Figure 1B; Methods). Performance across these external noise levels was modeled with the Perceptual Template Model (PTM) ^22^. The equivalent noise approach leverages a systematic manipulation of external noise to dissociate different sources of noise that limit perception. The method provides quantitative estimates of (i) baseline additive internal noise, (ii) stimulus-dependent internal noise that scales with stimulus strength, and (iii) the ability of the visual system to filter out external noise in the stimulus (unfiltered external noise), which is commonly associated with tuning of sensory neurons. A key insight behind the approach is that, at low levels of external noise, performance is primarily limited by baseline internal noise, and external noise only begins to affect performance at high noise levels that exceed baseline internal noise.

We estimated sensory noise associated with visual orientation perception at both group and individual levels. This allowed us to test whether individual differences in sensory noise predict individual variability in WM precision, and whether there is a group difference in sensory noise in SZ and NT. Importantly, we designed WM and noise estimation tasks to involve the same type of visual information, oriented visual stimuli, but to maximize differences in (a) stimulus salience, (b) task demands, and (c) how subjects report their responses. The orientation discrimination task focused on stimulus visibility, estimated contrast thresholds in noise (i.e., stimuli shown at low signal-to-noise levels), and required only a coarse categorical response with a keystroke (±45° left vs right tilt). In contrast, the WM task used highly visible noise-free grating stimuli (i.e., high signal-to-noise) and required a precise report of the remembered orientation with a dial. If we had measured perceptual noise using one feature, like orientation, and memory using another feature, like color, then differences in sensory noise between these features across individuals would be confounded with our measurements.

### High baseline internal noise is associated with worse WM performance in NT

We began by examining whether sensory noise measured from the orientation discrimination task was linked to WM performance in NT. In the visual WM task, NT exhibited the expected pattern of greater WM variance with increasing set size (Figure 1C, red symbols; *F*(2, 58) = 200.42, *p* < .001, *ηp²* = .87). In the orientation discrimination task, the same participants showed a highly characteristic nonlinear result, with contrast thresholds remaining fairly constant at low levels of external noise but increasing linearly (in log space) at higher levels of external noise (Figure 1D, red) ^22^. These data were modeled with the PTM to estimate the baseline internal noise, stimulus-dependent internal noise, and the unfiltered external noise for each participant (see Methods).

To test our hypothesis that sensory noise predicts WM capacity in NT adults, we used a multiple linear regression with the noise estimates from PTM as predictors of the average WM recall variance (Figure 2A & 2B, red). The model significantly accounted for 31.5% of the variance in WM precision (*F*(3, 26) = 3.99, *p* = .02, *R*^2^ = .32, *R*^2^*_Adj_* = .24). However, only the baseline internal noise and unfiltered external noise were significant predictors of WM recall variance on their own (baseline internal noise: *B* = 1.02, *SE* = .44, *t*(29) = 2.31, *p* = .03, *ηp²* = .17; unfiltered external noise: *B* = .21, *SE* = .09, *t*(29) = 2.31, *p* = .03, *ηp²* = .17). The stimulus-dependent internal noise (*B* = - 11.75, *SE* = 8.83, *t*(29) = -1.33, *p* = .20, *ηp²* = .06) did not significantly predict WM variance. This suggests that sensory noise associated with perceiving orientations is significantly associated with WM performance in NT adults.

**Figure 2.**
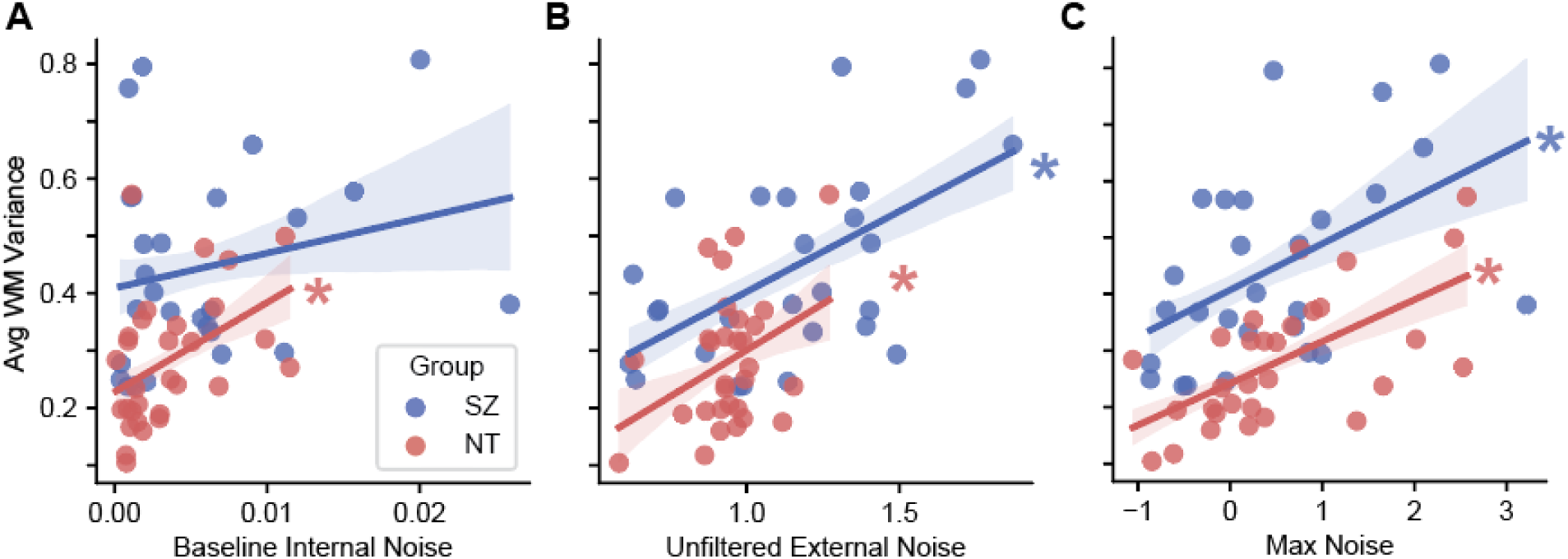
Sensory noise limiting orientation discrimination was associated with WM performance in both SZ (blue) and NT (red). **A**. In NT, baseline internal noise and unfiltered external noise were significant predictors of WM performance. **B**. In SZ, unfiltered external noise was a significant predictor of WM deficit. **C**. The ‘larger’ noise between the two (baseline internal noise vs. unfiltered external noise; z-scored per group) for each individual was significantly correlated with WM variance in both groups. This suggests that WM precision may be primarily constrained by the dominant source of sensory noise within an individual.

### Elevated sensory noise is associated with impaired WM performance in SZ

If sensory noise is a key limiting factor of visual WM precision, then a deficit in WM may be, at least in part, due to a deficit in sensory processing. We tested this hypothesis in SZ a disorder with a well-established WM deficit ^8^. In the WM task, individuals with SZ showed a marked impairment: the SZ group exhibited higher WM recall variance across all set sizes relative to NT (Figure 1C, blue; *F*(1,55) = 18.04, *p* < .001, *ηp²* = .25). There was also a significant group by set size interaction (*F*(2,110) = 5.05, *p* = .008, *ηp²* = .08) with individuals with SZ showing greater deficits at larger set sizes. Because the WM task relies on reporting remembered orientation with fine motor movements, we also conducted a control motor precision task where participants reproduced the orientation of visible gratings (see Methods). However, the results showed no significant differences between NT and SZ (*t*(55) = 1.65, *p* = 0.11, *ηp²* = .05), indicating that the observed WM impairment in SZ was not due to differences in motor precision.

In contrast to the well-established WM deficit in SZ, much less is known about visual orientation processing in SZ, particularly limitations related to sensory noise. Our results from the orientation discrimination task reveal a previously unknown perceptual deficit in SZ, with individuals with SZ exhibiting overall elevated contrast thresholds across a full range of external noise levels (Figure 1D, blue). These group differences were best explained by a model in which SZ and NT differed in the levels of both the baseline internal noise and unfiltered external noise. Specifically, the best-fitting model indicated a 61% increase in baseline internal noise along with a 21% increase in unfiltered external noise in the SZ group relative to the noise levels of the NT (R^2^ = .98). The model was significantly different from the null model which assumed that the two groups did not differ in their noise (*F*(2, 26) = 29.95, *p* < .001, *ηp²* = .70, ΔR² = .03), and was not significantly different from the full model where all three types of noise varied between the two groups (*F*(1,25) = .69, *p* = .41, *ηp²* = .03, ΔR² = .0004). These findings together suggest that SZ is associated with elevated levels of sensory noise.

Critically, similar to NT, atypically elevated levels of visual noise in the SZ group predicted their WM performance (Figure 2A & 2B, blue). To examine this relationship, a multiple regression analysis was carried out within SZ with individually estimated baseline internal noise and unfiltered external noise, shown to be different in SZ, as predictors. The analysis revealed that the baseline internal noise and unfiltered external noise together accounted for 34% of the variance in WM recall performance (*F*(2, 24) = 6.06, *p* = .007, *R*^2^ = .34, *R*^2^*_Adj_* = .28), with only unfiltered external noise significantly predicted WM recall variance on its own in SZ (*B* = .19, *SE* = .06, *t*(26) = 3.12, *p* = .005, *ηp²* = .29; baseline internal noise B = .29, SE = .44, *t*(26) = .67, *p* = .51, *ηp²* = .02).

Our results are consistent with the hypothesis that the noise associated with sensory processing limits visual WM. In NT, baseline internal noise and unfiltered external noise predicted individual differences in WM performance. In SZ, both baseline internal noise and unfiltered external noise were significantly elevated, with variations in the latter predicting variability in WM. One possibility is that WM precision is primarily constrained by the ‘larger’ noise (baseline internal noise vs. unfiltered external noise) that overrides the other within an individual, similar to the rationale for the equivalent noise paradigm explained above. To explore this further, we computed z-scores for both baseline internal and unfiltered external noise separately for the NT and SZ groups. Then, for each subject, we kept only the larger of the two z-scores. This simple max operation is intended to capture noise that is more prominent for that subject relative to that subject’s peer group (NT or SZ). We then correlated the result with WM performance and found that, for both NT and SZ, variability in sensory noise predicted visual WM precision (Figure 2C; NT: *r*(28) = .62, *p* = .0002; SZ: *r*(25) = .49, *p* = .009).

### Direct manipulation of sensory noise impacts WM performance in SZ

Next, we tested the observed link between sensory noise and WM in a causal design using transcranial direct current stimulation (tDCS) to affect the noise in the sensory processing (Study 2). The goal was to causally manipulate sensory noise and measure its influence on WM performance within SZ. Using tDCS involves the application of low-amplitude direct current to the scalp, thereby modulating the underlying excitability of cortical neurons in a polarity-specific manner. We focused on anodal stimulation because, in contrast to cathodal stimulation, it increases cortical excitability during and after stimulation, a modulation that is associated with increasing neural noise ^44–46^.

Twenty participants diagnosed with schizophrenia or schizoaffective disorder (SZ; Table 1) completed three stimulation conditions: anodal stimulation over occipitoparietal cortex, anodal stimulation over medial-frontal cortex, and sham (the location of which was randomly assigned to either occipitoparietal or medial-frontal cortex; Figure 3A). Stimulation over the occipitoparietal cortex served as our test condition based on prior work indicating that anodal stimulation over the visual cortex can heighten neuronal excitability underlying visual perception ^47^. Active stimulation over the medial-frontal cortex served as a spatial active control condition due to its distance from the target sensory regions. Following 20 minutes of either active 2.0 mA stimulation or 20 minutes of sham stimulation, participants immediately completed the same WM task and orientation discrimination task described in the experiments above.

**Figure 3.**
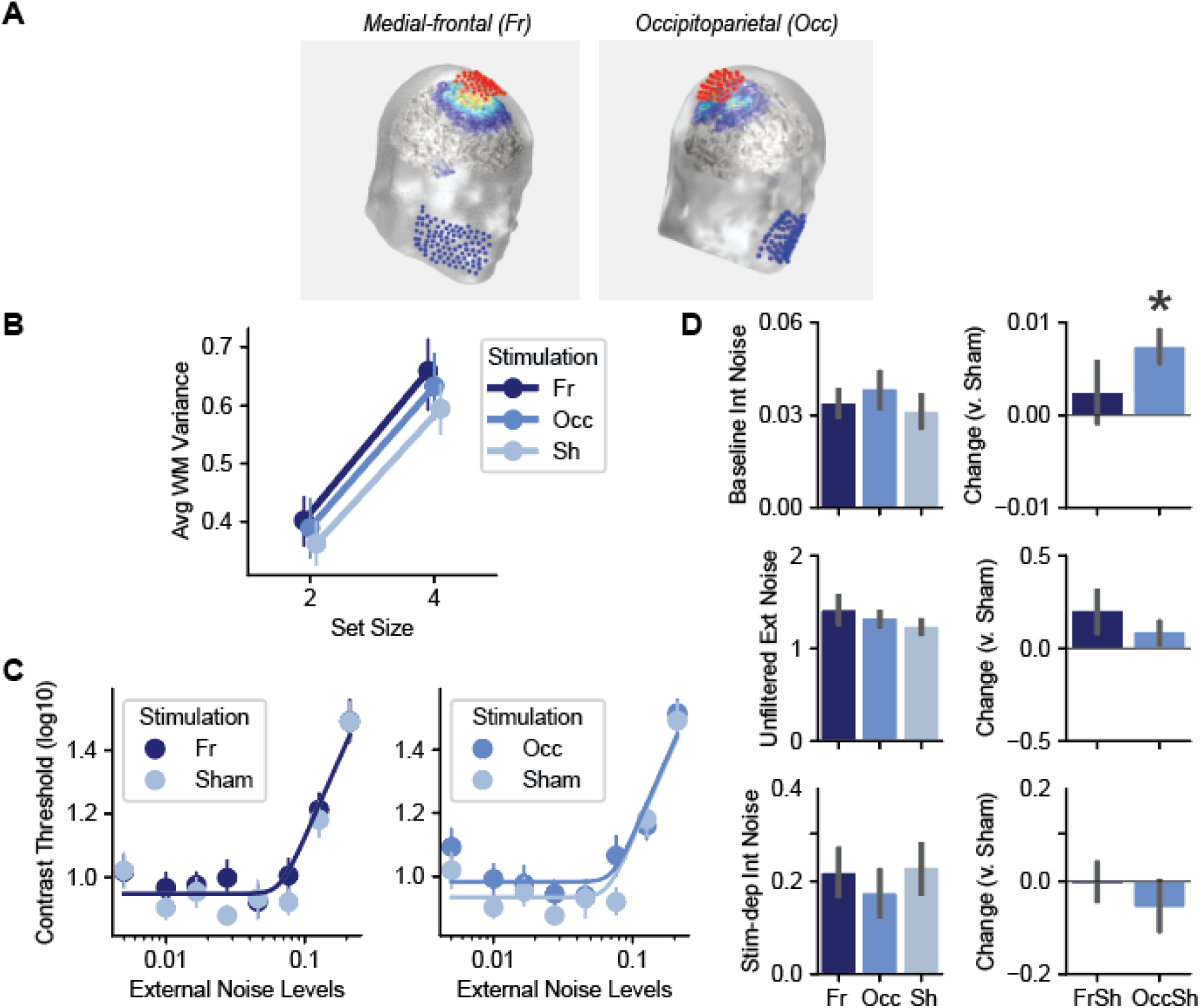
**A**. Modeled current distribution for the medial-frontal and occipitoparietal stimulation sites. **B**. Visual WM performance in SZ following tDCS across medial-frontal (Fr, darker blue), occipitoparietal (Occ, blue), and Sham conditions. (Sh, lighter blue), **C**. Visual orientation discrimination performance following tDCS. **D**. Changes in sensory noise estimated using PTM following tDCS. The parameter estimates from the hierarchical Bayesian model (Methods) are depicted for each stimulation conditions (left column) and for changes between active and sham conditions (right column; FrSh: medial-frontal vs. sham; OccSh: occipitoparietal vs. sham).

Across all analyses, the “change” following stimulation was defined as the difference between the active (Medial-frontal or Occipitoparietal) and sham stimulation conditions in either the WM or PTM noise metric.

We first tested whether stimulation over the occipitoparietal cortex would increase sensory noise, as measured by PTM noise parameters derived from the orientation discrimination task. Participants’ performance across the stimulation conditions again yielded the characteristic nonlinear dependency on external noise (Figure 3C). To characterize the effects of tDCS, the performances in the medial-frontal and occipitoparietal stimulation conditions were each compared with that in the sham condition separately using the PTM.

As hypothesized, visual orientation perception during the occipitoparietal stimulation was characterized by a significantly higher level of baseline internal noise compared to the sham condition (Figure 3D). Specifically, the effect of stimulation was best explained by a model in which active occipitoparietal and sham differed in the level of baseline internal noise. This model specified a 61% increase in internal noise with occipitoparietal stimulation relative to sham (R² = .96). The model fit was significantly better than a null model in which all noise parameters were constrained to be equal (F(1, 27) = 4.36, p = .046, ηp² = .14, ΔR² = .01), but did not differ from a full model in which all noise sources were allowed to differ from the sham condition (F(1, 27) = .42, p = .66, ηp² = .02, ΔR² = .001).

The increase in sensory noise was only observed following the occipitoparietal stimulation. Compared to the sham condition, we found no evidence that medial-frontal stimulation significantly impacted noise levels (Figure 3D). No model with any combination of varying noise parameters was significantly different from the null model in which all the noise parameters were held constant across active medial-frontal and sham conditions (all *F*s < 1.87, *p*s > .18; null model R^2^ = .97).

The observation that the anodal tDCS over the visual cortex increases sensory noise allowed us to test the hypothesized causal relationship between sensory processing and WM capacity. On average, similar to Study 1, SZ showed worsening WM performance with increasing set size (Figure 3B, *F*(1,19) = 61.52, *p* < .001, *ηp²* = .76), but, neither the medial-frontal nor the occipitoparietal tDCS stimulations significantly influenced the overall WM recall variance at the group level (main effect of stimulation condition: *F*(2,38) = 1.26, *p* = .29, *ηp²* = .06), with no significant interaction between set size and stimulation condition (*F*(2,38) = .17, *p* = .84, *ηp²* = .009).

However, the *concurrent changes* in sensory noise and WM performance *within individuals* were significantly correlated with each other following only the occipitoparietal stimulation (Figure 4B). In other words, although stimulation did not alter WM performance at the group level, individuals who showed a greater active-stimulation-related increase in baseline internal noise exhibited greater impairment in WM performance, but only following occipital stimulation. A multiple linear regression model with changes in all noise parameters (stimulation – sham) as predictors explained changes in WM performance (stimulation – sham), accounting for 45% of variance (*F*(3,14) = 3.79, *p* = .04, R^2^ = .45, *R^2^_Adj_* = .33), and only the changes in baseline internal noise on its own significantly predicted the changes in WM variance in these respective conditions (*B* = 12.56, SE = 5.31, *t*(17) = 2.36, *p* = .03, *ηp²* = .29; stimulus-dependent noise: B = -.30, SE = .15, *t*(17) = -2.05, *p* = .06, *ηp²* = .23; unfiltered external noise: *B* = .05, SE = .16, *t*(27) = .29, *p* = .78, *ηp²* = .006). In contrast, changes in the noise parameters following medial-frontal stimulation did not significantly explain changes in WM variance (Figure 4A; *F*(3,13) = 1.72, *p* = .21, *R*^2^ = .28, *R^2^_Adj_* = .12). Stated differently, increases in baseline internal noise specifically after occipitoparietal stimulation predicted the degree to which WM became less precise and more variable.

**Figure 4.**
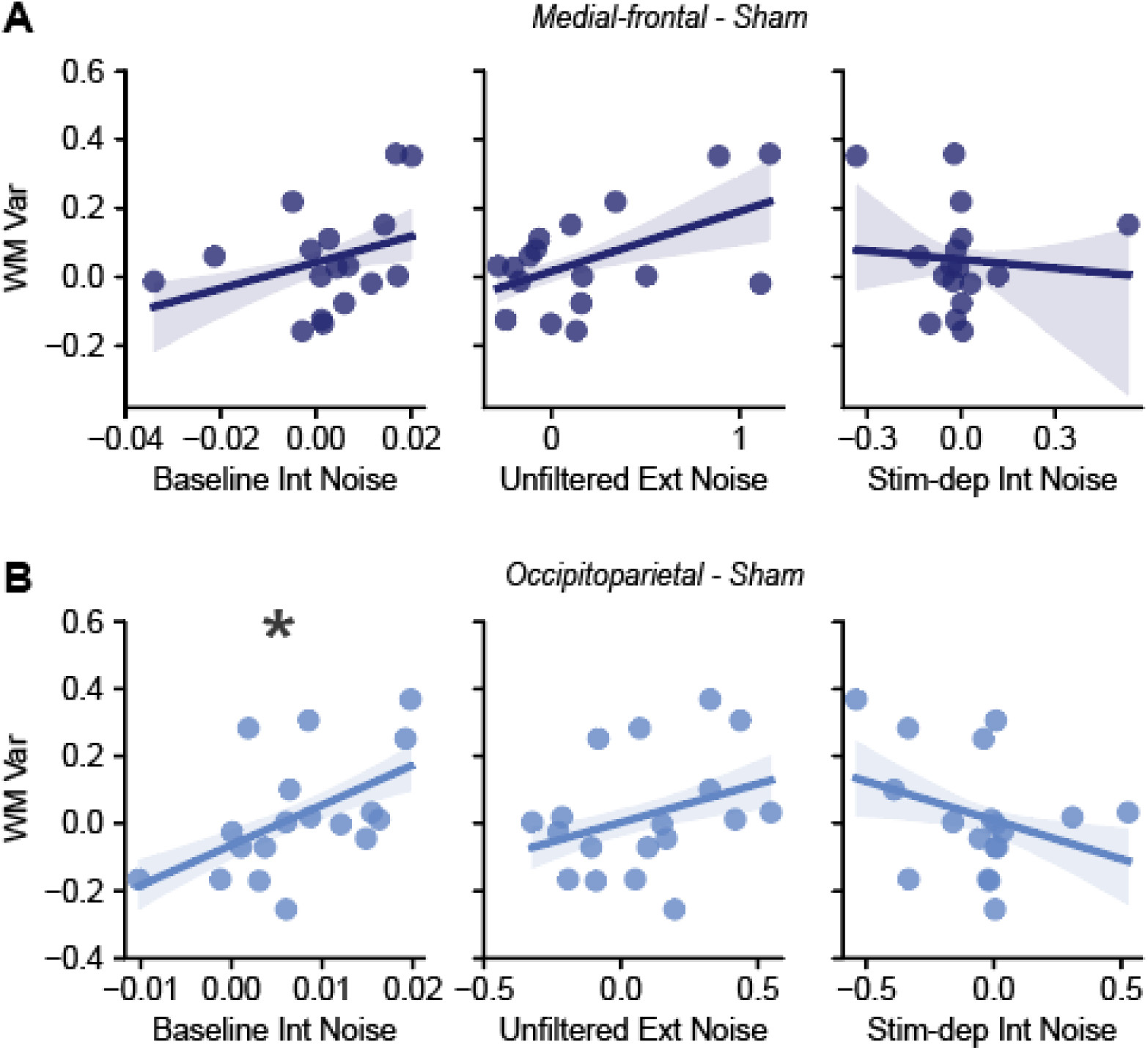
Relationship between changes in sensory noise and WM performance following tDCS stimulation in SZ. **A**. Changes in sensory noise following the medial-frontals stimulation did not significantly predict changes in WM performance as compared to sham. **B**. However, changes in baseline internal noise (left) following the occipitoparietal stimulation significantly predicted changes in WM performance.

### Modeling the effects of sensory noise on WM

To understand the mechanistic ways in which the sensory noise contributes to WM precision, we constructed a two-stage computational model of WM with encoding and maintenance stages (see Methods for details). The model draws from prior observations using delayed-estimation studies showing that WM precision degrades with set size and delay ^48^. A key assumption we make is that this reduction WM precision can arise from stochastic drift in memory representations during maintenance ^49^, with its rate determined by sensory noise.

Briefly, during encoding, stimulus orientations were represented by von Mises–tuned orientation channels with Poisson-like variability and decoded using a population-vector readout. Encoding precision decreased with set size. Sensory noise entered at this stage in two forms: unfiltered external noise broadened channel tuning, while baseline internal noise was modeled as circular Gaussian noise added to the decoded orientations.

During the maintenance stage (1s delay), memory precision degraded through “drift” in representations, implemented as diffusion of the encoded orientations over time ^49^ with faster diffusion at larger set sizes. Critically, we assumed that the same sensory noise that limits perception also adds variability to memory representations. Accordingly, the diffusion rate was multiplicatively scaled by a subject-specific sensory-noise factor, defined as the larger of the two z-scored noise sources estimated from the PTM. As a result, sensory noise reduced WM precision both by limiting encoding fidelity and by amplifying drift during maintenance.

We first simulated a full model in which PTM-derived baseline internal noise and unfiltered external noise could limit WM precision at three distinct stages as explained above (encoding tuning, post-decoding additive noise, and maintenance via diffusion during the delay period). Consistent with our observed empirical data (Figure 1C), the full model predicted reduced WM precision (higher WM variance) in the SZ group relative to the NT group (Figure 5A).

**Figure 5.**
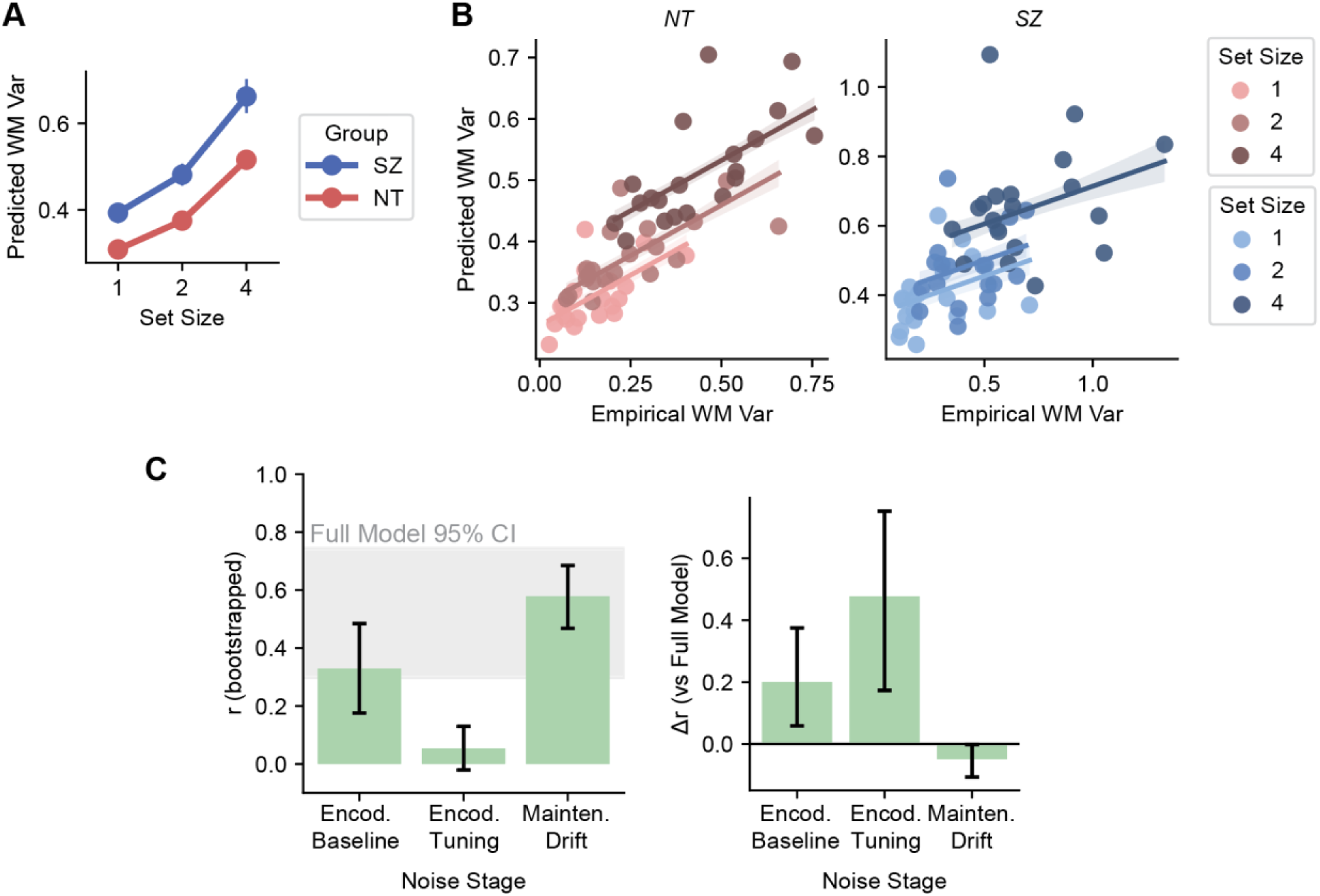
**A**. Group-averaged predicted WM variance as a function of set size for NT and SZ. **B**. Observed versus model-predicted WM variance for the full model across participants, shown separately for NT and SZ groups, with dots colored by set size and regression lines derived from mixed-effects models. **C**. Model comparisons. Left: bootstrapped Pearson correlation (r) between observed and predicted variance for models in which each noise component was separately simulated, with error bars showing 68% CI; the gray band indicates the 95% CI of the full model. Right: paired-bootstrap difference in correlation (Δr) relative to the full model, showing the relative contribution of baseline encoding noise, tuning-related encoding noise, and maintenance drift to explaining empirical observed performance. The error bars show 95% CI.

To quantify the relationship between empirical and model-predicted WM variance, we fit linear mixed-effects models in which the model-predicted WM variance was regressed on observed empirical WM variance, while accounting for repeated measurements across subjects and set sizes by including random intercepts for subject and for set size. Separate models were fit for the NT and SZ groups. In both groups, empirical WM variance significantly predicted model-predicted variance (Figure 5B; NT: β = .33, SE = .05, t(58) = 7.15, p < .001; SZ: β = .22, SE = .08, t(55) = 2.75, p = .008), indicating that the model explains inter-individual variability in WM precision.

To test whether this relationship between predicted and observed WM variance indeed depended on individual-specific sensory noise estimates, we conducted a permutation analysis in which the estimated PTM parameters were randomly shuffled across individuals within each group. Model goodness of fit was quantified as the summed partial R^2^ of the empirical WM variance predictor across groups, derived from the mixed-effects models similar to those above. The goodness-of-fit score obtained using the true subject-specific PTM estimates significantly exceeded the shuffled null distribution (permutation test, N = 1000; p = .01; true partial R^2^ = .59; shuffled mean partial R^2^ = .12, SD = .13), demonstrating that preserving individual sensory noise estimates significantly improved the model’s ability to explain inter-individual WM variability.

We then assessed which noise stage contributed most strongly to inter-individual variability in WM variance. For this, we ran reduced “ablation” simulations in which only one of the three noise stages was retained at a time (encoding tuning, post-decoding baseline noise, or maintenance drift). For each reduced model, we performed a bootstrap analysis with 1,000 iterations. Within each bootstrap iteration, subjects were resampled with replacement within group, and empirical and predicted WM variance were pooled across set sizes. Overall model goodness of fit was quantified as the Fisher-z-aggregated Pearson correlation between empirical and predicted variance across set sizes and groups.

The analysis revealed that sensory noise influencing the drift rate during the maintenance stage accounted for the largest share of inter-individual variability in WM variance (Figure 5C, left). To further quantify the relative contribution of each stage, we computed the difference in correlation coefficient (Δr) between the full model and each reduced model. Consistent with the ablation results, the model retaining only drift-related noise showed the most similar explanatory power to the full model (Δr closest to zero), suggesting that the sensory noise affecting the maintenance stage captured much of the variance explained by the full model (Figure 5C, right).

Finally, we asked whether the model could account for within-subject changes in WM variance following tDCS in the SZ group. Based on the tDCS and the above model results, we focused on a reduced model in which baseline internal noise modulates WM precision during the delay period by scaling the drift in the maintenance stage.

Accordingly, we simulated WM performance using a reduced model where only the maintenance-stage diffusion term was noise-dependent and made subject-specific predictions for sham and active stimulation conditions. For each participant, we obtained the mean WM variance across set sizes and computed the condition difference (stimulation − sham) for both empirical and predicted WM variance.

Consistent with our stimulation results, for occipitoparietal stimulation, changes in predicted WM variance were positively correlated with changes in empirical WM variance (r(16) = .50, p = .03). In contrast, the analogous relationship was not significant for medial-frontal stimulation (r(15) = .36, p = .15). Together, these results support the idea that the same sensory noise linked to the perception of visual orientations can constrain visual WM precision during maintenance.

## DISCUSSION

Our study provides a set of observations supporting the role of sensory noise in determining WM limits. The findings suggest that noise present in early stages of visual processing can limit the precision of WM at recall. Using a model of visual perception ^22,50^, we show that individual differences in sensory noise can both explain individual variability in WM capacity and account for the well-established WM deficit in individuals with schizophrenia. Our manipulations of sensory noise with tDCS over the visual cortex provide a causal link between sensory processing and WM, where changes in internal noise with stimulation resulted in corresponding changes in WM precision. Our computational model further suggests that the same sensory noise measured from visual perception can limit WM precision during maintenance by affecting the drift of WM representations. These findings indicate that lower-level sensory mechanisms contribute to capacity limitations typically attributed to high-level cognitive processes.

We conjecture that visual WM is particularly vulnerable to sensory noise not only because it relies on sensory information as its input, but more importantly, because it recruits sensory mechanisms for the maintenance of stored information ^17,21^. One putative advantage of involving sensory mechanisms for information maintenance in WM is the high fidelity of sensory processing. We posit that this is also what makes the precision of visual WM particularly sensitive to noise levels within the visual process.

Because effective WM performance depends on multiple stages, including encoding and maintenance ^51,52^, it is also susceptible to processing inefficiencies, including noise, at each of those stages ^4^. Both the sensory recruitment hypothesis and the sensory noise hypothesis suggest that sensory areas are actively recruited in these stages, where the fidelity of sensory representations directly influences WM precision. Our model suggests that sensory noise can also influence maintenance. Sensory noise likely is a potent factor even for maintenance because neural activity for memory retention in the sensory cortex is generally weaker than when a physical stimulus is present ^53^, leaving the representations more vulnerable to existing internal noise.

Our support for the idea that sensory noise affects WM storage does not preclude a role of the prefrontal circuitry in determining WM capacity ^18^. Instead, our findings add a complementary view from the sensory processing perspective: in tasks that rely on sensory details, such as orientation, color, or location, the fidelity of sensory cortex representations is crucial. Studies have shown that WM capacity limits can be observed even when visual stimuli remain in view ^54^. This suggests that WM capacity results not simply from remembering and maintaining representations but from performing simultaneous tasks on multiple representations. In this context, sensory noise is critical as it constrains the precision with which representations can be manipulated and compared. At the same time, cognitive processes also shape operations to be performed on these representations ^55^. Thus, WM capacity is a joint outcome of limitations in representational fidelity and the efficiency of higher-order control processes.

Our results also show that the WM deficits in SZ, traditionally associated with impairment in prefrontal processing, can be explained by elevated levels of sensory noise. Our observation is consistent with a recent finding that implied faster WM decay in SZ ^56^, which could result from noisy sensory representations. At the group level, the WM precision in SZ was best explained by their ability to filter out external noise. In the PTM, external noise filtering occurs at the earliest stages of visual processing, where ‘templates’ selectively enhance relevant signals and suppress irrelevant external noise. The effectiveness of this filtering depends on the tuning properties of these templates. Notably, prior work has demonstrated broader orientation tuning in SZ ^57^, which could explain elevated levels of unfiltered external noise observed in our study. It is possible that noisy representations due to broader orientation tuning in SZ are having a larger effect on WM capacity than elevated baseline internal noise.

We also show that direct manipulation of baseline internal noise through brain stimulation affects WM capacity in SZ. Anodal tDCS is known to increase cortical excitability ^58,59^, which can modulate the spontaneous fluctuation in neural activity. Given the lack of temporal precision and task-specificity in tDCS (stimulation is continuously applied for 20 minutes prior to completing the tasks), it is less likely that the stimulation selectively enhances task-related activity. If the tDCS rather amplifies random neural variability, it can reduce the overall signal-to-noise ratio in neural responses by increasing baseline neural noise. Our findings are not at odds with the previous observation of increased visual sensitivity to low-contrast stimuli following anodal tDCS ^60^, as in our task, increased excitability would amplify the responses for not only the target signal (oriented grating) but also the external noise added to the stimuli. It is also worth noting that the effects we observed were individual-specific: individuals with SZ who showed an increase in baseline internal noise following tDCS to the occipitoparietal cortex also exhibited a corresponding decline in WM precision. Recognizing this variability provides critical considerations for designing individualized intervention strategies.

Our findings identify the early visual system as a significant WM bottleneck in SZ. Elevated sensory noise in visual cortex has far-reaching consequences in addition to contributing to WM deficits. Other aspects of visual information processing may also be impacted by increased sensory noise in SZ, including motion perception ^35^ and visual context processing ^61^. Furthermore, face discrimination deficits associated with increased internal noise in SZ ^36^ could potentially cascade to social cognition impairments. Increased sensory noise has also been observed in individuals with autism spectrum disorders (ASD) ^43,62^, a population similarly noted for their WM deficits ^63,64^. In both SZ and ASD, elevated sensory noise may reflect or contribute to an excitation/inhibition imbalance in cortical circuits ^34^, a mechanism that has been widely proposed in both populations ^65–67^. In other words, increased sensory noise may play a mechanistic role in shaping higher-level cognitive outcomes such as WM in these populations. Understanding how sensory noise interacts with developmental and neuroanatomical factors, as well as its potential interactions with medications, could provide key insights into the shared and distinct pathways through which WM deficits emerge in a wide range of neuropsychiatric disorders ^6,51,68–71^.

In conclusion, we report both the correlational and causal evidence for the effects of sensory noise on WM capacity. We conclude that the noise that limits the efficiency in perceptual processing plays a critical role in constraining WM recall precision. The results offer new insights into the sensory recruitment hypothesis of WM and highlight sensory noise as a possible target for intervention (e.g., through brain stimulation or perceptual training) that could enhance WM capacity in populations whose WM deficits have been primarily associated with cognitive dysfunctions.

## METHODS

### PARTICIPANTS

Demographic and clinical information for participants who completed Studies 1 and 2 is summarized in Table 1. Twenty-seven (48% women; Study 1) and 20 (45% women; Study 2) medicated and clinically stable outpatients with chronic schizophrenia or schizoaffective disorder (SZ) were recruited from outpatient facilities in Nashville for participation. Diagnoses were made or confirmed according to the Diagnostic and Statistical Manual of Mental Disorders, Fourth Edition, Text Revision (DSM-IV-TR) criteria using the Structured Clinical Interview for DSM-IV (SCID-I/P) ^72^. SCIDs were administered by masters-level clinical psychology graduate students.

In Study 1, 30 (47% women) neurotypical controls (NT) were recruited from the same metropolitan area through advertisements. NT had no history of DSM-IV Axis I disorders or family history of psychosis. We estimated premorbid IQ using the North American Adult Reading Test (NART) ^73^. SZ and NT in Study 1 were matched on age, estimated IQ, and handedness but not on years of education (*t*(55) = 4.18, *p* < .01). Exclusion criteria included a history of head injury, neurological disorder, or substance abuse in the 6 months preceding the study.

All SZ participants were taking medications at the time of the study; in Study 1, 26 were taking atypical antipsychotics, 1 was taking a typical antipsychotic, 4 were taking mood stabilizers, and 10 were taking SSRIs, and in Study 2, 19 were taking atypical antipsychotics, 1 was taking a typical antipsychotic, 1 was taking a mood stabilizer, and 10 were taking SSRIs. Antipsychotic doses were converted to chlorpromazine equivalents for comparative analyses. Symptom severity in SZ was assessed with the Brief Psychiatric Rating Scale (BPRS; Overall and Gorman, 1962), the Scale for the Assessment of Negative Symptoms (SANS) ^74^, and the Scale for the Assessment of Positive Symptoms (SAPS) ^75^. All participants were screened for normal or corrected-to-normal vision with the Snellen test of visual acuity. For the Snellen test, participants read through the chart using one eye at a time, viewed the letters from a distance of 20 feet, and read letters from the top of the chart aloud to an experimenter who scored their performance. Based on this viewing distance and the reference standard (20/20), the 8^th^ row of letters from the top of the chart consisted of letters subtending an angle of 5° with each letter part subtending 1°.

All participants provided written informed consent to study procedures approved by the Vanderbilt University Institutional Review Board and were compensated for their participation at a rate of $20 per hour.

### STIMULI, TASKS, AND PROCEDURE

All tasks and stimuli in Studies 1 and 2 were programmed in MATLAB and Psychtoolbox ^76,77^ and presented on a linearized monitor (20-inch Sony CRT; 1024 x 640 resolution; 120 Hz) on a gray background. A chin rest was used to maintain viewing distance at 77 cm with each pixel subtending 0.036°. An experimenter was present in the room during both visual discrimination and working memory tasks to encourage task engagement. Task order was counterbalanced across participants.

#### Visual Working Memory Precision Task

To assess visual WM, participants completed a WM task in which they were instructed to remember locations and orientations of sinewave gratings (1 cycle/deg) in a two-dimensional raised cosine envelope (radius = 1°). All gratings were presented at maximum contrast to ensure visibility. The trial procedure was as follows: first, a memory array consisting of one, two, or four oriented gratings was presented for 2 s. Orientations were randomly selected from all possible options [-90°, 90°] with the constraint that orientations within a given memory array were unique. Grating locations for each trial were randomly chosen from eight positions around an invisible circle (radius 4°) centered on a fixation point. After a short delay (1 s), a new, randomly oriented [-90°, 90°] grating appeared at one of the prior locations (test probe). Participants used an input dial (PowerMate USB Mulitmedia Controller; Griffin Technology) to adjust the test probe’s orientation so that it matched the orientation of the grating presented at that same location. Participants were asked to be as precise as possible when matching orientations and responses were untimed. Participants indicated their final response by pressing down on the dial, leading to a 3 s inter-trial interval. Trials began with a centrally-presented dynamic fixation sequence ^78^. Participants completed a total of 450 trials, split equally into 10 blocks with rest periods lasting a minimum of 30 s.

Because group differences in ability to use the manual dial may confound hypothesized group differences in WM precision, participants also completed a short motor precision task that did not tax memory. Participants completed the motor precision task prior to the WM task in order to become acclimated to using the manual dial. Trials involved presentation of a randomly oriented grating located at one of eight positions around an invisible circle (radius 4°) that remained on the display. A randomly oriented grating was then presented at the center of the display, and participants were instructed to use the manual dial to adjust the orientation of this central grating until it was aligned with the orientation of the first grating, which remained on the display until the participant’s response was submitted. Consistent with instructions on the WM task, participants were asked to be as precise as possible when matching orientations and responses were untimed. Participants completed a total of 24 trials such that each of the 8 possible grating locations was sampled three times.

This manual recall method has been used with both healthy adults and psychiatric populations to obtain a more detailed, continuous measure of visual WM precision compared to typical binary match/non-match response paradigms ^21,79^. Importantly, this method enables WM performance to be analyzed as a distribution of recall errors (the difference between input response and target orientation) with accompanying descriptive properties of the distribution. As the number of visual features or items to be remembered increases, the decline in recall precision is captured by the increasing variance of the error distribution.

#### Orientation Discrimination Task

Participants performed a coarse orientation discrimination task ^43^ where they judged the tilt of centrally presented gratings with the same spatial frequency and size as those in the WM task (tilted ±45° from vertical). Gratings were temporally embedded in two independent external noise samples. The grey-level pixel values for external noise were sampled from a Gaussian distribution with a mean of 0 and SD that varied across trials, such that its root mean square contrast defined the external noise level to be one of 8 set levels ranging from 0-21% (0, 0.3, 0.61, 1.24, 2.51, 5.1, 10.3, or 21%). The rapid temporal succession of noise, stimulus, and noise frames (16.7 ms duration per frame) results in an intentional perceptual merging of the grating and noise frames that is consistent with PTM experiments, as it allows for finer adjustments of stimulus contrast ^50^. Participants indicated their response by a left or right arrow key press, and brief auditory feedback (50 ms tone) was provided after correct trials to maintain task engagement. As in WM task, trials started with a dynamic, centrally-presented fixation sequence. Participants completed 680 trials, split equally into 8 blocks with rest periods lasting a minimum of 30 s.

The stimulus contrast in the visual discrimination task was adjusted every trial using the Functional Adaptive Sequential Testing (FAST toolbox ^80^) procedure, which determines the most informative contrast levels to be sampled for estimating participants’ contrast thresholds. Two FAST structures were interleaved to estimate psychophysical functions reflecting the contrast level required to elicit reliable orientation discrimination at two different sensitivity levels (d’=1.089 and 1.634), corresponding to 70.71% and 79.37% accuracy. Of note, data analyses were completed separately from the within-task FAST procedure in order to limit biased threshold estimations from potential increased accidental error responses in SZ. See Park et al. (2017) ^43^ for a complete description of the psychophysical procedure.

### TRANSCRANIAL DIRECT CURRENT STIMULATION

The tDCS was administered (Study 2) using a battery driven, constant current stimulator (MindAlive Inc., Alberta, Canada) and a pair of conductive rubber electrodes (active: 19.25 cm2, reference: 52 cm2). The electrodes were placed over saline-soaked sponges and each held in place with elastic cloth headbands. In the active anodal conditions, current is applied for 20 minutes. Stimulation intensity is fixed at 2.0 mA, as this intensity demonstrated improvement in visual acuity in a prior study ^47^. For medial frontal stimulation, the anodal electrode was placed over the medial frontal lobe at midline (site FCz of the International 10-20 System) and the cathodal electrode as placed over the right cheek to avoid confounding effects from other brain regions. For occipitoparietal stimulation, the anodal electrode was placed over occipitoparietal cortex at midline (site Pz of the International 10-20 System), with the cathodal electrode again placed over the right cheek. Consistent with tDCS montages in related work, the placement of the cheek electrode is on a diagonal, parallel to the jaw line, 3 cm from the lip corner (cheilion) above the jaw ^47,81^. Location of anodal electrode placement over FCz or Pz was determined with individual scalp measurements. We modeled the current flow using COMETS and our previously described methods for estimating the fields generated by the stimulator ^59,82^.

The tDCS sham condition involves similar administration as that for the active condition, except the stimulation only lasts 30 seconds, which ramps up and down at the beginning and end of the 20-minute period. This sham procedure results in the same physical sensations (e.g., tingling, itching) that are reported with active tDCS ^47^. Following each session, participants completed a questionnaire and visual analog scale that included questions regarding attention, concentration, mood, vision, headache, fatigue, and skin sensations during tDCS. Participants were additionally asked to rate whether they had received active versus sham stimulation and the degree of confidence associated with those ratings. Participants tended to overestimate the presence of stimulation, leading to an accuracy rate of detecting the correct stimulation condition of 57%. However, the hit rate for detecting sham stimulation was 15%, well below chance, suggesting that participants were blind to the sham versus active conditions. Additionally, confidence ratings did not differ across stimulus condition (p > .23). Scores across all questionnaire ratings did not significantly differ between stimulation conditions (ps > .092).

The present study followed a within-subjects design, such that each participant completed the three stimulation conditions (anodal active over medial-frontal, anodal active over occipitoparietal, and sham) on three separate days. Session order was counterbalanced across participants, and the location of sham (medial frontal/occipitoparietal) was randomly assigned. The time interval between session visits was at least 48 hours to minimize potential task practice effects or carryover effects related to the previous session’s brain stimulation (M=6.60 days between sessions, SD=4.11 days) ^83^.

All study sessions began with 20 minutes of active or sham tDCS applied over medial frontal or occipitoparietal cortex. Immediately after tDCS, participants completed the visual WM precision task (20-30 minutes), followed by the visual discrimination task (40-60 minutes). For the WM task, participants completed a shortened version of the visual WM task described above in order to reduce task duration and cognitive burden on participants. Participants completed a total of 60 trials, split equally into 3 blocks with rest periods lasting a minimum of 30 s. While the trial procedure was identical, memory arrays only consisted of two of four oriented gratings. In total, the two tasks lasted approximately 90 minutes, which is within the time window in which tDCS effects have been shown to last in comparable stimulation procedures ^81^.

### ANALYSES

#### Visual Working Memory Task Analysis

Grating orientations were recorded and reported in the parameter space of all possible line orientation values [-90°, 90°) and converted to the circular space [-π, π) radians. Recall error on each trial was calculated as the difference between the orientation of the target grating and the orientation of the test probe that was reported (input with the manual dial) by the participant. Given that internal noise is hypothesized to add variability to, or destabilize, the internal representation ^21,39^, the primary performance metric of interest on the visual WM task was participants’ recall variance. Poorer WM performance is thus reflected by higher recall variance, as demonstrated by the characteristic increase in recall variance (decline in precision) as the number of items remembered increases ^37,38^. The Von Mises probability density function (circular normal distribution) with mean *θ* = 0 (no error, such that recall is centered at the target memory location) and *σ* was used to fit every participant’s distribution of recall errors at each memory array set size. The Von Mises probability density function for a given angle *x*, centered at *θ* with a concentration *κ* where *I*_0_(*k*) is the modified Bessel function of order 0:

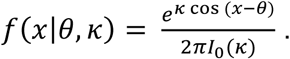

The variance of the Von Mises probability density function (*σ*^2^), which reflects the dispersion of the distribution, is analogous to the inverse of the distribution’s concentration (1/*κ*). The concentration parameter (*κ*) was estimated from each participant’s recall errors at every memory array set size using a maximum likelihood method. Estimates of distribution concentration were converted to variances by taking the inverse for subsequent between-group and individual differences analyses.

#### Orientation Discrimination Task

Data from the visual discrimination task were analyzed with two different approaches. The first involved a conventional PTM analysis to estimate model parameters at a group level, thus allowing us to test for relative groups differences in levels of noise in the three noise parameters between SZ and NT. The second approach utilized hierarchical Bayesian model fitting to estimate each individual’s model parameters. The latter thus allowed for a better understanding of individual differences in noise estimates. It also enabled us to examine how noise estimates related to WM performance.

#### Conventional PTM Analysis

To examine group-level differences in noise estimates, visual discrimination performance was first analyzed using a conventional PTM fitting method. First, psychophysical contrast thresholds were estimated for each participant at each of the 8 external noise levels with data pooled across the two FAST structures. Individuals’ contrast thresholds for orientation discrimination were estimated with Weibull functions at every external noise level using the following equation:

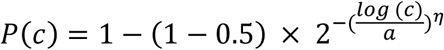

where *P* represents accuracy (percent correct), *c* is stimulus contrast, *α* is threshold at 75% accuracy, and *η* is the slope. The two free parameters, *α* and *η*, were estimated with a Bayesian model fitting method implementing a Markov Chain Monte Carlo (MCMC) technique. Group averages of individuals’ estimated thresholds were used to fit the conventional PTM. Coefficient indices for each noise source (unfiltered external noise (*A*_*e*_), baseline internal noise (*A*_*a*_), stimulus-dependent internal noise (*A*_*m*_)), were introduced into the standard PTM to account for differences in noise levels between groups:

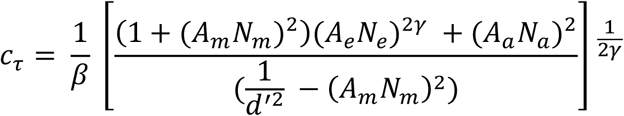

During PTM fitting, these three SZ coefficients (*A*_*e*_, *A*_*a*_, *A*_*m*_) could freely vary or be fixed at 1, with the latter constraining a given noise estimate to be equal across groups (Study 1) or stimulation conditions (Study 2) and the former allowing for relative differences between groups or stimulation conditions in a given noise estimate. To determine if noise sources differed between groups or conditions, all possible candidate models accounting for each combination of free and fixed noise estimates were run: the null model, which assumed no differences in any of the noise sources, the full model, which assumed differences in all three noise sources, and six remaining models reflecting the different possible combinations of the three noise estimates. In addition to which ever noise coefficients were specified to freely vary, all candidate models included the following four free parameters: stimulus-dependent internal noise (*N*_*m*_), internal additive noise (*N*_*a*_), signal gain from the perceptual template (*β*), and the exponent of the nonlinear transducer function (*γ*). Thus, the full model included 7 free parameters, the null model included 4 free parameters, and the other six models included between 5 and 6 free parameters. The eight candidate models were each fitted to average thresholds using a least-squares method. Candidate models were compared by evaluating differences in goodness-of-fit using *r*^2^. The best fitting model was determined with the F-test for nested models. Specifically, the best fitting model was identified as the model with the fewest free parameters that was not significantly different from the full model, in which all parameters could freely vary.

#### Hierarchical Bayesian Model Analysis

A hierarchical Bayesian modeling technique was used to fit each participant’s data with the PTM in order to obtain individual noise parameter estimates and examine relations between noise estimates and visual WM performance ^43^. Unlike in the conventional PTM analysis approach described above, in which model parameters are estimated from the contrast thresholds at each level of external noise, the hierarchical Bayesian approach allows for model parameters to be estimated for each participant. The following PTM equation was used:

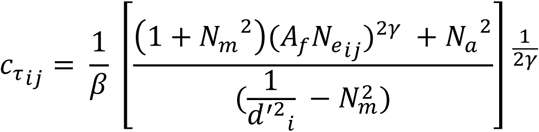

where the contrast threshold is estimated at a given accuracy criterion, *i* (either 70.71% or 79.37%), and external noise level, *j*. Each participant’s response on a given trial (with a given external noise level and at a given difficulty level) was assumed to be drawn from a Bernoulli distribution to capture the set of possible outcomes from the 2-alternative forced-choice response in the visual discrimination task. PTM fitting was constrained such that the gain parameter *β* and nonlinear power parameter *γ* were held constant to minimize the number of free model parameters while enabling estimation of all three noise parameters of interest (*N*_*a*_, *N*_*m*_, *A*_*f*_) and the slope of the psychometric function (*η*). Values were set at *β* = 1 and *γ* = 2.64 for Study 1, and *β* = 1 and *γ* = 1.5 for Study 2, for all individuals, based on estimates generated from the conventional PTM analysis. Importantly, modifications in the values of the gain parameter *β* and nonlinear power parameter *γ* did not change the results regarding relations between noise estimates and WM recall variance. For Study 1, based on results from the conventional PTM analysis which showed a group difference in baseline internal noise and unfiltered external noise, only the *N*_*a*_ and *A*_*f*_ noise parameters were set to vary across individuals. Free model parameters were estimated for each individual with the MCMC method for sampling from posterior probability distributions. A given individual’s model parameters were constrained by hierarchical priors in that parameters were assumed to be drawn from a group’s (SZ or NT) population distribution with their own means and SDs. Priors for the two groups were set to broad uniform distributions.

#### Statistics

Repeated measures ANOVAs were conducted to determine changes in recall variance with increasing WM set size and to probe group differences (Study1) and effects of tDCS locations (Study 2). To assess the relationship between the sensory noise estimates and WM variance, multiple linear regressions were completed within each group. Nested F-tests were used to compare different models of PTM with varying numbers of free parameters. Statistical analyses were performed at the significance level of .05.

### COMPUTATIONAL MODELING OF WORKING MEMORY PRECISION

#### Model

The model consists of two stages: encoding of stimulus orientations by a population code and maintenance of the encoded representation via diffusion during a delay. Subject-specific sensory noise parameters estimated from the PTM were used to modulate encoding tuning, post-decoding noise, and memory drift.

#### Encoding stage

On each trial, a stimulus orientation *θ* is encoded by a population of *M* orientation-tuned neurons with preferred orientations 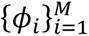. The mean response of a neuron *i* is given by a von Mises tuning function:

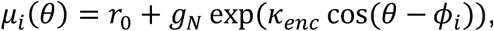

where *r*_0_ is the baseline firing rate, *g*_*N*_ = *g*_0_/*N* is the gain scaled by set size *N*, and *k*_*enc*_ is the orientation tuning. Unfiltered external noise *n*_*ext*_ estimated from the PTM modulates encoding precision through setting the sharpness of the tuning curves:

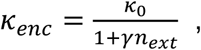

where *k*_0_is the baseline tuning sharpness, and *γ* controls the influence of the unfiltered external noise. Neural responses included Poisson-like variability, approximated as Gaussian noise with variance proportional to the mean response:

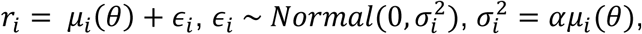

where *α* controls the strength of the variability. Following orientation decoding, baseline internal noise was added as a form of additive circular Gaussian noise applied to the decoded orientation:

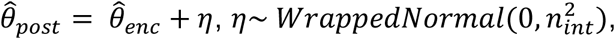

where *n*_*int*_ is mapped from the PTM baseline internal noise estimate.

#### Maintenance stage

During the delay period of duration *T*, the memory representation undergoes diffusion on the circular orientation space. Memory noise is modeled as a Gaussian random walk with variance increasing linearly over time:

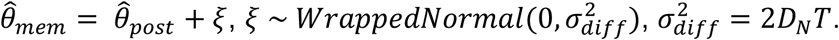

The diffusion constant *D*_*N*_ depends on set size and subject-specific sensory noise:

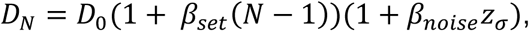

Where *D*_0_ is the baseline diffusion rate, *β*_*set*_ controls the effect of set size, and *z*_*σ*_ is the dominant internal noise defined as the larger of the z-scored baseline internal noise and unfiltered external noise.

#### Behavioral error and precision

Memory error was defined as the circular difference between the remembered and true orientations. Memory precision was quantified by the concentration of the error distribution across trials, converted to an equivalent von Mises concentration parameter. WM variance was reported as the inverse of this concentration (1/κ).

Subject-specific PTM estimates included baseline internal noise and unfiltered external noise. These parameters were z-scored across subjects and mapped linearly onto model parameters governing three noise stages: (i) orientation tuning during encoding, (ii) post-decoding noise, and (iii) diffusion strength during maintenance. Mapping coefficients were calibrated by fitting the model to match the mean empirical WM variance across set sizes. All parameters unrelated to sensory noise or diffusion were held constant across simulations and serve only to define the scale of the population code.

#### Analyses

##### Relationship between predicted vs empirical WM variance

For each subject and model variant, we simulated 1,000 trials per condition to predict WM precision using a full model in which PTM-specific sensory noise affected WM precision at all three noise stages. Model predictions were generated separately for each subject and set size.

To assess whether the model captured individual differences in WM performance, we related empirical and model-predicted WM variance using linear mixed-effects models. Model-predicted WM variance was regressed on empirical WM variance while accounting for repeated measurements across subjects and set sizes via random intercepts for subject and set size. Models were fit separately for NT and SZ.

##### Permutation test of individual specificity

To test whether individual-specific sensory noise estimates were critical for model performance, we conducted a permutation analysis in which PTM parameters were randomly shuffled across individuals within each group. For each permutation, model predictions were recomputed and related to empirical WM variance using the same mixed-effects framework. Model goodness of fit was quantified as the summed partial R^2^ of the empirical predictor across groups, derived from the mixed-effects models. The observed goodness-of-fit score was compared against the null distribution to obtain a permutation p-value.

##### Bootstrap ablation analysis

To determine which noise stage contributed most to inter-individual variability in WM precision, we performed a bootstrap analysis on the reduced model variants. In the reduced models, sensory noise was allowed to affect only one stage at a time, with other noise parameters fixed to group-median values. In each bootstrap replicate (1,000 iterations), subjects were resampled with replacement within group, and empirical and predicted WM variance were pooled across set sizes. Overall model goodness of fit was quantified as the Fisher z–aggregated Pearson correlation between empirical and predicted WM variance. Confidence intervals were obtained from the bootstrap distribution, and differences in correlation relative to the full model (Δr) were computed to assess similarity in explanatory power.

##### tDCS analysis

To examine whether the model could account for changes in WM performance following tDCS in SZ participants, we focused on a reduced model in which baseline internal noise modulated diffusion during the maintenance stage. WM variance was simulated for sham and stimulation conditions, and condition differences (stimulation − sham) were computed for both empirical and predicted WM variance, averaged across set sizes. Pearson correlations were used to assess the relationship between changes in predicted and empirical WM variance for occipito-parietal and frontal stimulation sites.

## DATA AVAILABILITY

The data generated and/or analyzed during the current study are not publicly available due to participant confidentiality concerns. Although the data do not contain direct identifiers, they include information that, in combination (e.g., location, patient group), may pose a risk of re-identification. De-identified data will be made available from the corresponding authors on request, subject to institutional review and approval.

## CODE AVAILABILITY

Code to run experiments and analyze data (MATLAB and Python) are available on Github repository: https://github.com/gt-core-lab/wm-noise-sz.

## ACKNOWLEDGMENTS

Supported by the NIH (R00-EY034546 to WJP; T32-EY007135, P30-EY08126 to GFW; R01-MH110378 to GFW and SP, R01-MH073028, R01-MH128967 to SP), Smithgall Watts Early Career Award to WJP, Brain & Behavior Research Foundation (NARSAD Independent Investigator Grant to DT; Distinguished Investigator Grant to SP), NSF (BCS-2147064 to GFW), the E. Bronson Ingram Family (to GFW), American Psychological Foundation (Alexander Gralnick Research Investigator Award to SP), and Gertrude Conaway Vanderbilt Endowment (to SP).

## AUTHOR CONTRIBUTIONS

Conceptualization: WJP, MI, GFW, DT, and SP; Methodology: WJP, MI, GFW, DT, and SP; Software: WJP, and MI; Formal analysis: WJP, GH, and MI; Investigation: WJP, and MI.; Writing—Original draft: MI and WJP; Writing—Review & Editing: WJP, GFW, DT, and SP; Funding acquisition: GFW, DT, and SP.

